# Large ribosomal subunit, eIF5B, Met-tRNA_i_^Met^ and mRNA cooperate to complete accurate initiation

**DOI:** 10.1101/2020.05.30.125153

**Authors:** Jinfan Wang, Jing Wang, Byung-Sik Shin, Thomas E. Dever, Joseph D. Puglisi, Israel S. Fernández

## Abstract

Recognition of a start codon by the first aminoacyl-tRNA (Met-tRNA_i_^Met^) determines the reading frame of messenger RNA (mRNA) translation by the ribosome. In eukaryotes, the GTPase eIF5B collaborates in the correct positioning of Met-tRNA_i_^Met^ on the ribosome in the later stages of translation initiation, gating entrance into elongation. Leveraging the long residence time of eIF5B on the ribosome recently identified by single-molecule fluorescence measurements, we determined the cryoEM structure of the naturally long-lived ribosome complex with eIF5B and Met-tRNA_i_^Met^ immediately before transition into elongation. The structure uncovered an unexpected, eukaryotic specific and dynamic fidelity checkpoint implemented by eIF5B in concert with components of the large ribosomal subunit.

**One sentence summary:** CryoEM structure of a naturally long-lived translation initiation intermediate with Met-tRNA_i_^Met^ and eIF5B post GTP hydrolysis.

---

Protein synthesis starts with the assembly of a ribosomal complex at the start site on the mRNA (*1*). Eukaryotes employ numerous translation initiation factors (eIFs) to achieve the 80S initiation complex (IC) with the initiator aminoacyl-tRNA (Met-tRNA_i_^Met^) and the AUG start codon of the mRNA programmed in the ribosomal peptidyl-tRNA site (P site) (*2*). The 40S small ribosomal subunit, accompanied by eIFs and Met-tRNA_i_^Met^, is recruited to the 5’-UnTranslated Region (5’-UTR) of the 7-methylguanosine-capped mRNA (*3*). This is followed by a dynamic inspection of the 5’-UTR in search of the correct AUG codon as a start site, a process termed “scanning” (*4, 5*). Upon AUG recognition by the Met-tRNA_i_^Met^ anticodon, a series of eIF reorganizations and departures coupled to ribosomal conformational rearrangements occurs, resulting in a post-scanning 48S preinitiation complex (PIC) (*6–8*). This 48S PIC is then joined by the 60S large ribosomal subunit to form the 80S IC, catalyzed by the universally conserved GTPase eIF5B (Fig. 1A) (*9, 10*). Using single-molecule fluorescence methods, we recently revealed that maturation of the *Saccharomyces cerevisiae* 80S IC to the elongation-competent state (80S EC) with an exposed codon in the A site for aminoacyl-tRNA delivery is gated by the slow eIF5B dissociation from the complex (Fig. S1, A and B) (*11*). This dissociation of eIF5B requires GTP hydrolysis, with the timing of its dissociation critical to stringent start codon selection.

**Fig. 1.**
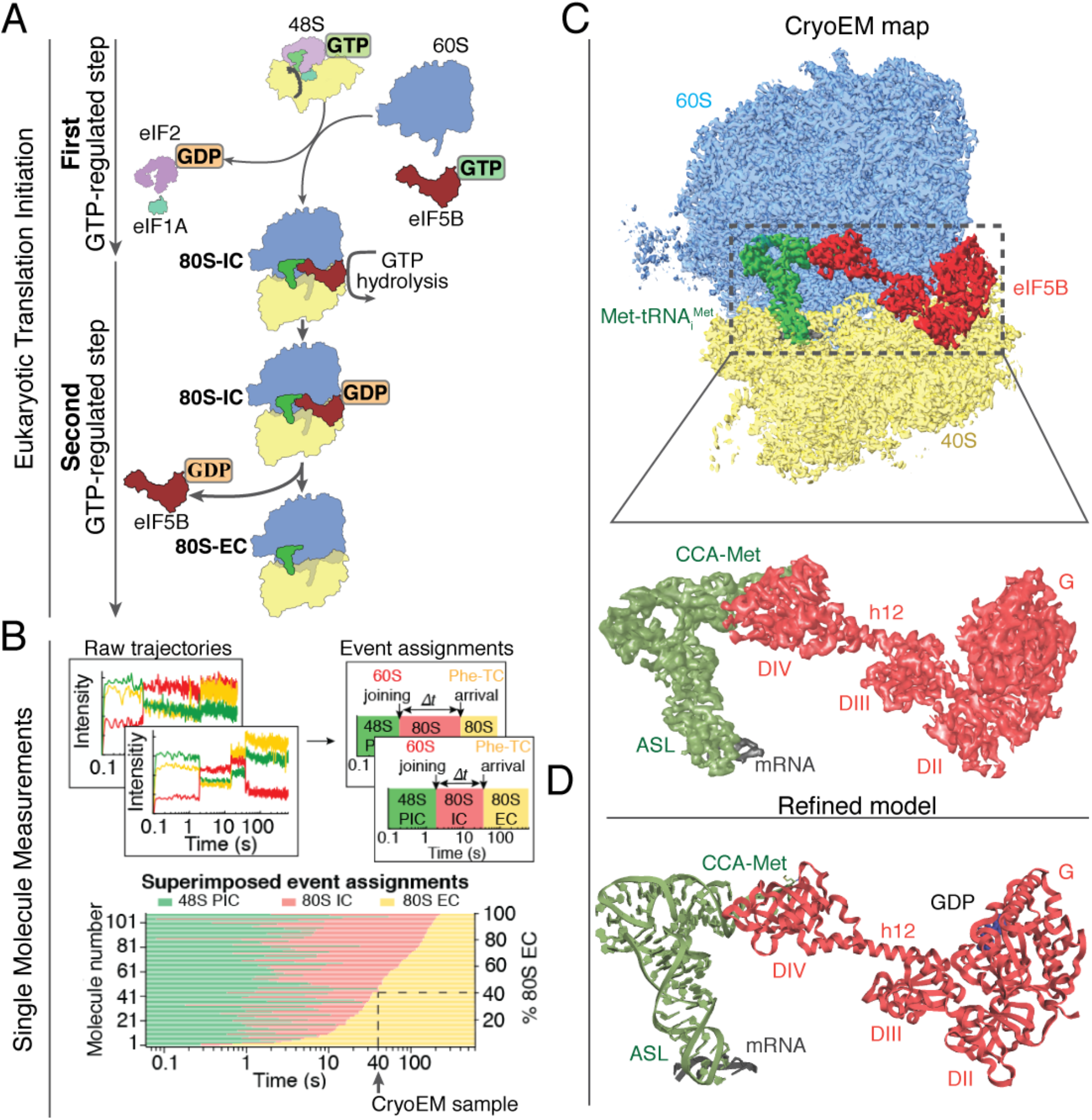
CryoEM sample preparation guided by single-molecule fluorescence data and overall architecture. **(A)** Overview of translation initiation in eukaryotes. Two steps, regulated by GTPases eIF2 and eIF5B control the delivery of Met-tRNA_i_^Met^. **(B)** CryoEM samples were prepared and froze at the timepoint 40 s after mixing 48S PICs, eIF5B:GTP and 60S. At this timepoint, ~40% of the total 80S particles are expected to be 80S EC, while the other ~60% are 80S IC with eIF5B-bound, as demonstrated by our previous single-molecule fluorescence data (also see Fig. S1) (*11*). **(C)** Overview of the final cryoEM map obtained after maximum likelihood classifications in Relion (*29*). Large subunit (60S) is colored in blue, small subunit (40S) yellow, Met-tRNA_i_^Met^ green, mRNA grey and eIF5B red. Bottom, details of the local map obtained for Met-tRNA_i_^Met^, mRNA and eIF5B. Domain IV (DIV) of eIF5B is in close contact with the acceptor stem of Met-tRNA_i_^Met^. **(D)** Stereochemically refined models for Met-tRNA_i_^Met^, mRNA and eIF5B with components indicated.

These dynamics results left unanswered whether the slow dissociation of eIF5B was limited by its GTPase-activation/GTP-hydrolysis, Pi release, and/or eIF5B:GDP dissociation. Previous cryoEM reconstructions have described the architectures of the 80S IC bound with eIF5B:GDPCP at medium resolution, representing snapshots of the assembly prior to eIF5B GTP hydrolysis (*12, 13*). In addition, Met-tRNA_i_^Met^ has sequence features that allow it to bind directly to the P site of the ribosome during initiation, unlike elongator tRNAs (*14–16*). Here, guided by the kinetics determined by our previous single-molecule fluorescence measurements, we performed cryoEM analysis of the on-pathway initiation complexes to provide high-resolution information on the molecular mechanism by which eIF5B escorts the initiator tRNA into the ribosomal P site and gates the transition from initiation to elongation.

Leveraging the slow dissociation of eIF5B from the 80S IC in the native initiation pathway (average lifetime of the eIF5B-bound 80S state is 30-60 s at 20°C) (*11*), we prepared and froze samples at a pre-steady-state reaction timepoint (40 s, Fig. 1B, Fig. S1 and Methods) corresponding to when ~60% of the assembled 80S should be bound with eIF5B (80S IC) after mixing 48S PICs with eIF5B:GTP and 60S. Image processing and 3D classification in Relion3 (*17, 18*) identified a homogeneous class of ~70% of the total pool of 80S particles, allowing the reconstruction of a 3D map with a global resolution of 2.9 Å (Fig. S2, S3 and S4). The reconstruction shows clear density for a tRNA in the surroundings of the P site and density for all domains of eIF5B (Fig. 1C). A further analysis of the map revealed excellent densities for domain III (DIII), the connecting α-helix 12 (Fig. 1C and D, h12 (*19*)) and especially for domain IV (DIV) of eIF5B, as well as for the Met-tRNA_i_^Met^ and the start-codon of the mRNA (Fig. 1C and D and Fig. S4). Less well resolved were domains II and G of eIF5B and the elbow region of Met-tRNA_i_^Met^ (Fig. S3).

In our reconstruction, DIV of eIF5B is tightly packed against the peptidyl transferase center (PTC) of the large subunit where it contacts the _73_ACCA_76_-Met end of Met-tRNA_i_^Met^ (Fig. 2). The close interaction of eIF5B DIV with the acceptor stem of Met-tRNA_i_^Met^ enforces a change in the trajectory of the _73_ACCA_76_-Met end when compared with the position adopted by the acceptor stem on an elongation peptidyl-tRNA in a canonical configuration (Movie S1) (*20*). This distortion is not limited to the _73_ACCA_76_-Met of tRNA_i_^Met^, as the whole acceptor stem is distorted compared with its position in an elongation tRNA (Fig. 2B and C). The cluster of G-C base pairs of the acceptor stem specific to Met-tRNA_i_^Met^ seems to play a pivotal role in this context, allowing a specific distortion of this stem as the Met-tRNA_i_^Met^ simultaneously interacts with eIF5B and the start-codon (Fig. 2A and B and Fig. S5). Basic residues of the domain IV of eIF5B make specific interactions with this G-C base pairs cluster (*21*); Arg955 of eIF5B contacts the base of G70 of Met-tRNA_i_^Met^ from the major groove of the acceptor stem (Fig. 2E and F). Although mutating Arg955 to Ala in eIF5B did not substantially alter the growth rate of yeast in rich medium, the mutant strain could not grow under amino acid starvation conditions (Fig. 2G), suggesting impaired *GCN4* expression that could result from ribosomes scanning past the start codon (termed “leaky scanning”) of the stimulatory first upstream open reading frame (uORF1) in the mRNA. Thus, the interaction between eIF5B Arg955 and Met-tRNA_i_^Met^ G70 may also play an important role in start-site selection. Consistently, deletion of eIF5B DIV has also been shown to enhance leaky scanning in yeast (*19*).

**Fig. 2.**
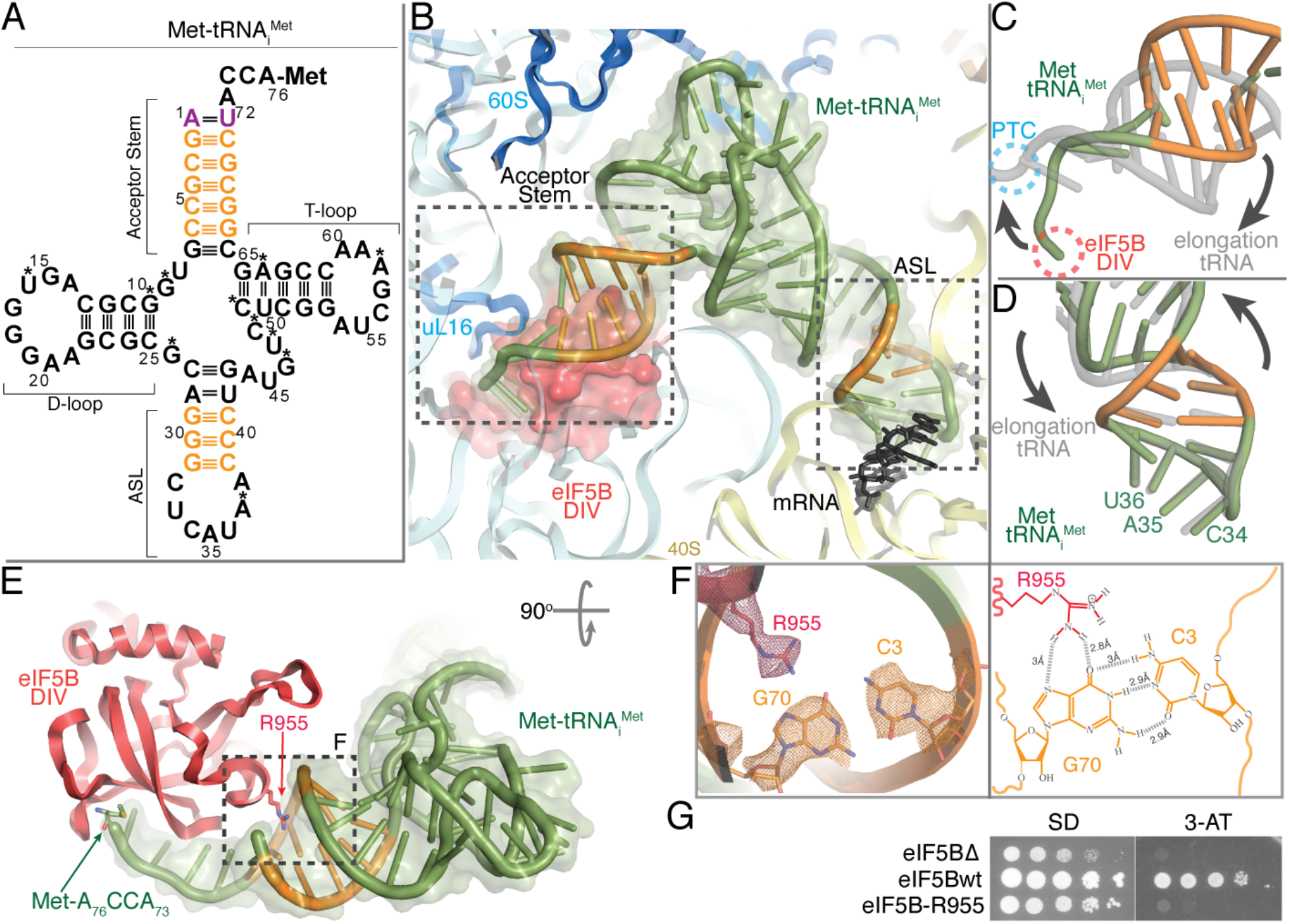
Role of the G-C base pairs clusters in Met-tRNA_i_^Met^ conformation. **(A)** Sequence diagram of the *S. cerevisiae* Met-tRNA_i_^Met^ with initiator-specific base pairs colored in orange and modified nucleotides marked with asterisks. **(B)** Overview of the conformation adopted by Met-tRNA_i_^Met^ on the 80S-IC complex with the initiator-specific G-C base pairs colored in orange. **(C)** Superposition of Met-tRNA_i_^Met^ (green) with a P site classical-state elongation tRNA (grey) reveals a distorted configuration of the acceptor stem of Met-tRNA_i_^Met^ when compared with an elongation peptidyl-tRNA. This distortion prevents the _74_CCA_76_-Met end of Met-tRNA_i_^Met^ from reaching the peptidyl transferase center (PTC, blue) on the 60S. **(D)** In contrast, the ASL of Met-tRNA_i_^Met^ has reached it final position when compared with an elongation tRNA in the P site. **(E)** The arginine residue 955 at DIV of eIF5B specifically recognizes the Hoogsteen edge of the base of nucleotide G70 of Met-tRNA_i_^Met^ (red arrow, R955). **(F)** CryoEM density for side chains and individual bases in the area around eIF5B R955. On the right, chemical diagram with contacts and distances indicated. (**G**) Yeast growth assay of strains lacking eIF5B or expressing wild-type or R955A mutant spotted on minimal (SD) or starvation (3-AT) medium; impaired growth on 3-AT medium points towards a defect in translation start-site selection on the *GCN4* mRNA in the mutant strain.

In contrast, the anticodon arm of Met-tRNA_i_^Met^ features a configuration very similar to that described for a peptidyl-tRNA in an elongation, canonical state (Fig. 2D, Fig. S5B and Movie S1) (*20*). Anticodon bases C34, A35 and U36 have reached their final elongation position, which would allow a productive transition into elongation. The rigidity contributed by the G-C cluster of base-pairs in the ASL seems to play an important role, allowing a local distortion that guarantees an ideal positioning of the anticodon bases to maximize the interaction with the AUG codon (*15*), while allowing the tRNA to bend at the elbow region of T-loop/D-loop tertiary interaction to prevent accommodation of the aminoacyl-acceptor stem to the elongation state (Fig. 2B and C). Thus, both G-C base pair clusters at the acceptor stem and the anticodon arm of Met-tRNA_i_^Met^ are essential to allow a specific conformation of the Met-tRNA_i_^Met^ on the P site in both the early (*21*) and late stages of initiation (*14*).

For the mRNA, we could unambiguously identify six nucleotides including the AUG start codon and the three nucleotides immediately upstream (Fig. 3). Only weak densities ascribable to the A site codon could be observed, and no ordered RNA density could be identified at the entry nor exit sites of the mRNA channel on the 40S. This is in marked contrast with 48S PIC structures from earlier initiation stages, in which long stretches of the mRNA from the entry to the exit sites were well resolved (*5*), pinpointing a less prominent role of mRNA/40S interactions once initiation has progressed into its later stages.

**Fig. 3.**
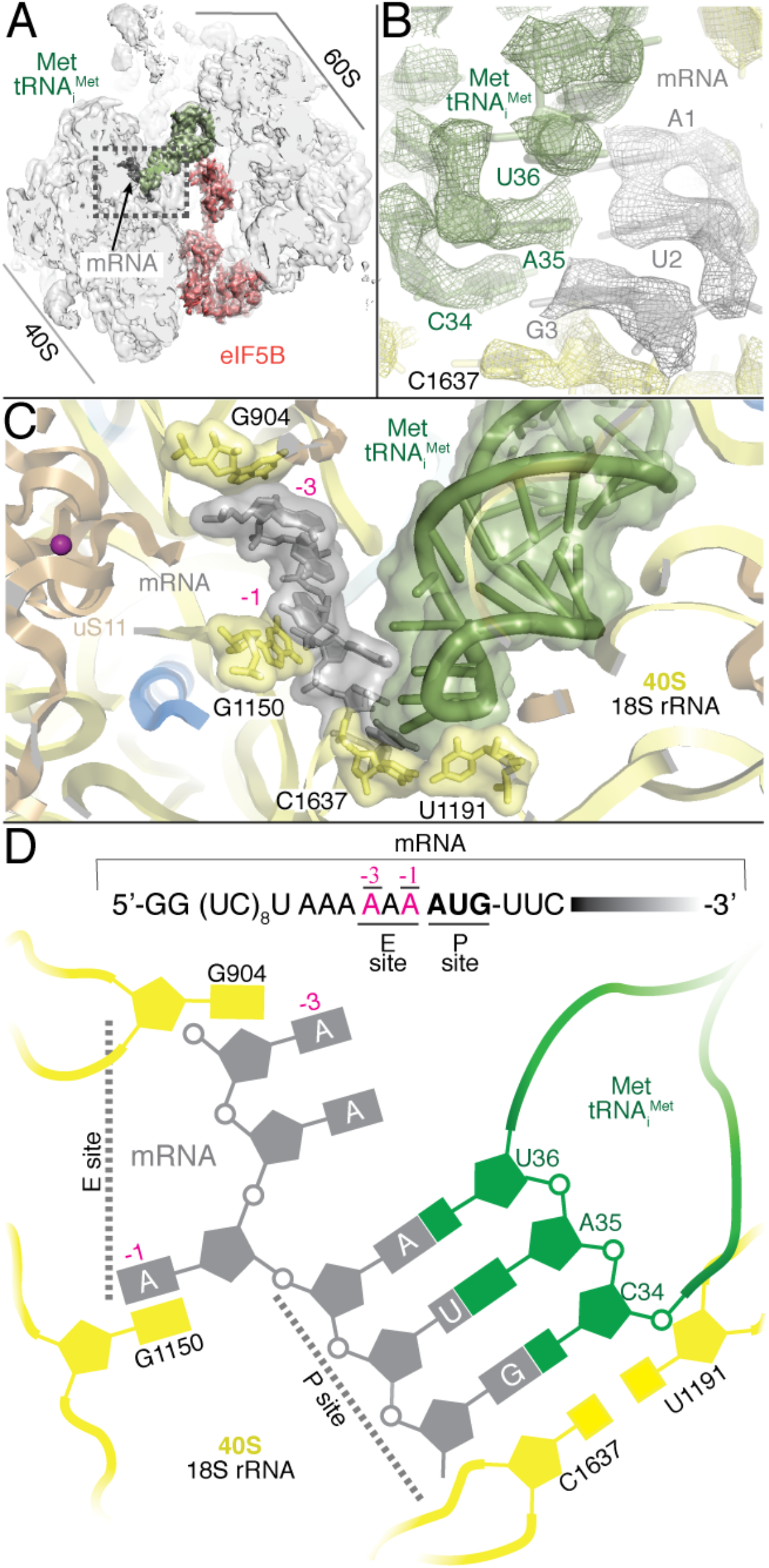
mRNA nucleotides at positions −1/−3 play a key role in later stages of initiation. **(A)** Overall view of the 80S/Met-tRNA_i_^Met^/eIF5B complex in an orientation centered on the mRNA. **(B)** Detailed view of the small ribosomal subunit P site with experimental cryoEM density shown. The Met-tRNA_i_^Met^ ASL is in green, the start-codon grey and ribosomal bases of the small subunit yellow. **(C)** Overall view of the P and E sites of the small ribosomal subunit. Six bases corresponding to the start codon of the mRNA and three preceding bases corresponding to nucleotides −1 to −3 showed unambiguous densities and could be modeled (grey). The final refined model with labeled components are shown. **(D)** Simplified schematic view of the conformation adopted by the start codon in the P site and the preceding −1 to −3 bases in the E site in the presence of eIF5B.

The codon-anticodon interaction observed in our reconstruction is virtually identical to a canonical cognate pair (Fig. 3B and C) (*20*). Nucleotides C1637 and U1191 of the 18S rRNA at the base of the 40S P site bracket the base pair mRNA-_3_G:C_34_-tRNA_i_^Met^ in a very similar configuration as in an elongation complex (Fig. 3C and D). Additionally, ribosomal bases G1150 and G904 engage mRNA nucleotides at position −1 and −3, respectively, in stacking interactions in a similar configuration as in early initiation complexes in the 48S PIC context (Fig. 3C and D) (*6, 7*). Thus, in the later stages of initiation, prior to entrance into elongation, the start-codon and its flanking residues present a hybrid configuration with bases at position −1/−3 retaining key contacts with rRNA bases that are instrumental in the early positioning of the mRNA on the 40S and, at the same time, with the start-codon features characteristic of a standard conformation of an elongation state (*20*). Hence, the start codon-surrounding sequence, especially the bases at positions −3/−1, plays an essential role along all initiation, from early “scanning” to later entrance into elongation (*22*).

The high quality of the map around the PTC region allowed a precise modeling, revealing specific contacts of residues of eIF5B DIV with the four terminal bases of the Met-tRNA_i_^Met^ as well as with ribose and phosphate backbone atoms (Fig. 4). Specifically, the base of Met-tRNA_i_^Met^ nucleotide A76 is narrowly monitored by eIF5B residues Glu921 and His924, which anchors the adenine moiety to eIF5B DIV (Fig. 4B and C). In this orientation, the methionyl group esterified to the 3’OH of the A76 ribose is directed towards a hydrophobic “pocket” formed by the surface of eIF5B around residue Ile874 (Fig. 4C and D and Fig. S6).

**Fig. 4.**
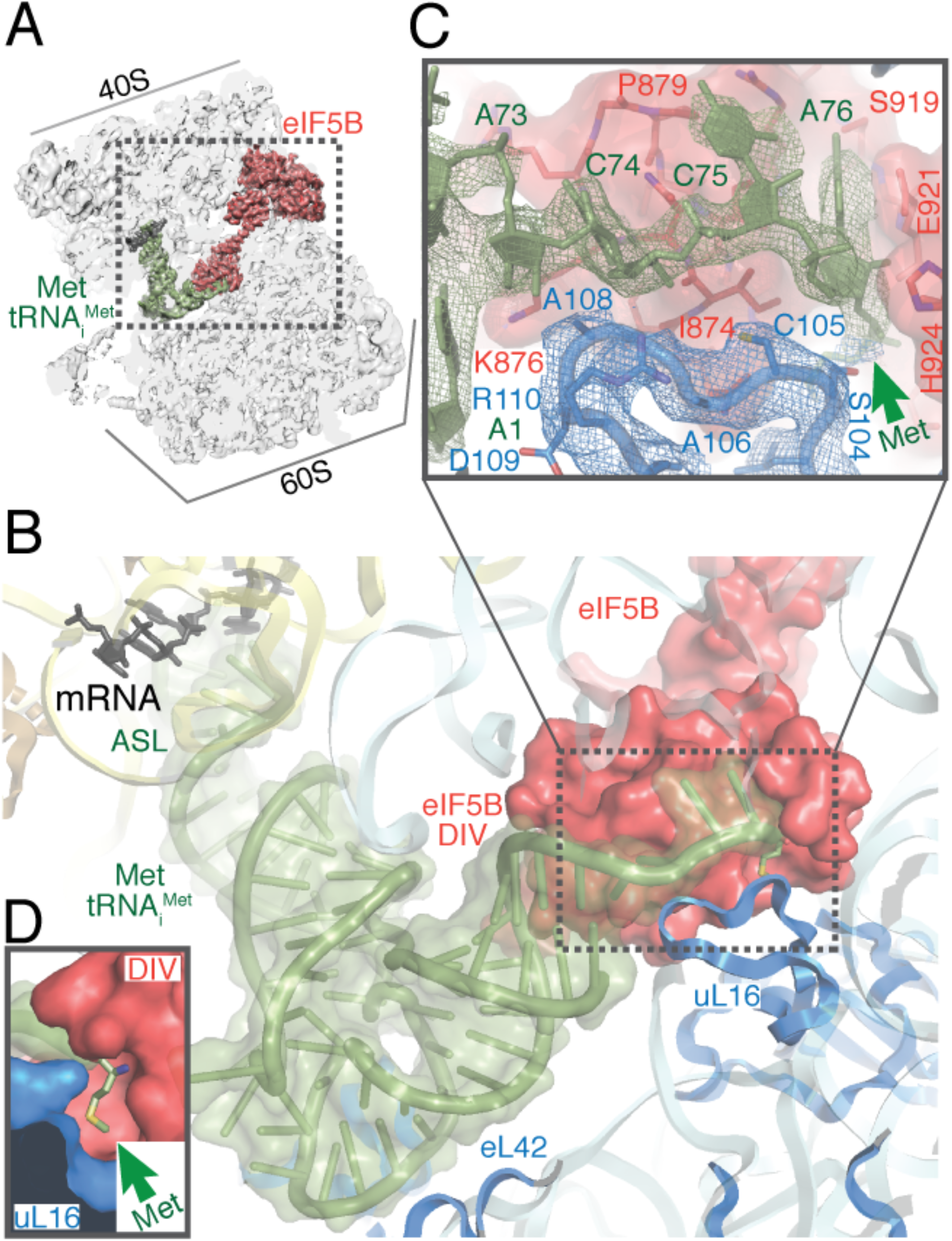
Ribosomal protein uL16 collaborates with eIF5B DIV in Met-tRNA_i_^Met^ A76-Met check. **(A)** Overall view of the 80S/Met-tRNA_i_^Met^/eIF5B complex in an orientation centered on eIF5B DIV. **(B)** A detailed view of the Met-tRNA_i_^Met^ conformation adopted while simultaneously interacts with the mRNA and eIF5B DIV on the 80S. **(C)** A detailed view of eIF5B DIV focused on the acceptor stem region of Met-tRNA_i_^Met^. DIV of eIF5B is shown in red as semi-transparent Van der Waals surface, Met-tRNA_i_^Met^ nucleotides depicted in green with experimental cryoEM density shown and uL16 residues are in blue with experimental cryoEM density shown. DIV of eIF5B and residues 102-110 of uL16 form a hydrophobic cavity where the methionine residue of Met-tRNA_i_^Met^ is hosted, as shown in **(D)**.

Intriguingly, this “hydrophobic pocket” is capped by a loop of ribosomal protein uL16, a component of the 60S (Fig. 4B-D and Fig. S6), which is conserved in yeast and humans. This uL16 loop, formed by residues 100 to 120, is disordered in reported elongation 80S complexes, but is well resolved in our reconstruction, allowing its modelling and refinement. Notably, this loop is stringently checked at the last step of 60S biogenesis by a sophisticated cellular machinery that blocks 60S exporting to the cytoplasm if its integrity is compromised (*23*). No clear function has been assigned for this loop that would justify such a conserved and costly cellular machinery (*24*).

Here, residues Ser104 to Arg110 of this loop are located in close proximity to the _73_ACCA_76_-Met of Met-tRNA_i_^Met^, in an almost parallel configuration to the phosphate backbone of the Met-tRNA_i_^Met^ (Fig. 4C). Additionally, uL16 residues Leu103 and Ser104 tightly approach the eIF5B residues around Leu871 to define a narrow and hydrophobic cavity where the methionine residue is hosted (Fig. 4D and Fig. S6). No obvious contacts between the methionine moiety and either eIF5B or uL16 residues could be observed, which points towards an overall chemical requirement for an amino acid attached to an initiator tRNA in terms of size and hydrophobicity rather than a specific amino acid identity. Indeed, when tRNA_i_^Met^ is mis-acylated with glycine, Gly-tRNA_i_^Met^ is active in 48S PIC and 80S formation (Fig. S7). However, subunit joining rate is ~9-fold slower and the subsequent transition to elongation became ~3-fold faster. Both effects might reflect weakened interactions between tRNA_i_^Met^ and eIF5B/uL16 when the methionine moiety was replaced by the smaller, non-hydrophobic glycine residue.

Analysis of the G domain of eIF5B that binds GTP reveals clear density for a bound nucleotide that however lacks features compatible with the presence of a γ-phosphate, implying that the nucleotide state is either GDP or GDP-Pi (*25, 26*). Moreover, switch I of eIF5B is disordered, which differs from the ordered state prior to GTP hydrolysis (*25, 27*). Thus, our reconstruction represents an intermediate of the 80S IC right before its transition to the elongation-competent state, but post GTP hydrolysis. This is further supported by the fact that the ribosomal inter-subunit configuration observed here is different from the 80S IC state prior to GTP hydrolysis (Fig. S8). A ~3° counterclockwise rotation of the 40S in relation to the 60S was observed in the pre-GTP hydrolysis state, which was coupled with apparent 40S head swivel (Fig. S8 (*12, 13, 28*)). However, the eIF5B-bound 80S IC observed here presents a configuration very similar to a canonical non-rotated 80S complex, with virtually no rotation of the small subunit and minimal swiveling of the 40S head (Fig. S8). Thus, the 80S IC will reconfigure its conformation after GTP hydrolysis to a state that is more similar to the non-rotated elongation-competent state, and this reconfiguration is coupled to the structural rearrangements of eIF5B.

In summary, our structural analysis of the pre-steady-state initiation complexes identified a novel intermediate of the late translation initiation complex on the native reaction pathway. Our results demonstrated that GTP hydrolysis is not the rate-limiting step for eIF5B dissociation from the 80S IC, but the subsequent Pi release and/or eIF5B-GDP dissociation. Notably, our structure describes stable interactions among Met-tRNA_i_^Met^, eIF5B DIV and uL16 after GTP hydrolysis by eIF5B. Disruption of this network of contacts is required for release of eIF5B, perhaps explaining the slow eIF5B departure rate post GTP hydrolysis. Such cooperation among Met-tRNA_i_^Met^, an initiation factor and the 60S subunit highlights a eukaryote-specific mechanism to control the progression of the initiation complex into the elongation phase.

## Acknowledgments

We are grateful to Dr. Dong-Hua Chen for technical support for sample freezing, and the cryoEM facility staff at Columbia University Irving Medical Center (CUIMC) for excellent technical assistance in cryoEM data acquisition. This work was supported by the US National Institutes of Health (NIH) grants GM011378 and AI047365 to J.D.P.; a Knut and Alice Wallenberg Foundation postdoctoral scholarship to J.W. (KAW 2015.0406); the Intramural Research Program of the NIH (T.E.D).

## Supplementary Materials

### Materials and Methods

#### Materials

All the yeast *Saccharomyces cerevisiae* 40S and 60S ribosomal subunits, initiation factors eIF1, 1A, 2, 5, and 5B, and mRNA were prepared and characterized as described (*1*). The model mRNA-consensus, with the sequence GG(UC)_8_U<u>AAAAAA</u>AUGUUCAAAUAA(UC)_16_, was an uncapped, unstructured model mRNA containing the optimal yeast 5’ context consensus sequence (underlined) with a biotin covalently linked to the 3’end (*1*). Native yeast methionylated initiator tRNA (Met-tRNA_i_^Met^) was purchased from tRNA Probes, LLC (MI-60).

#### 80S:eIF5B complex assembly on model mRNA

Previously by applying single-molecule fluorescence microscopy methods to a purified, reconstituted yeast translation system, we have revealed that eIF5B is the gating factor during the transition from eukaryotic translation initiation to elongation (*1*). We preformed post-scanning 48S preinitiation complexes (48S PICs, wherein 40S was labeled with a Cy3 dye) which were immobilized on zero-mode waveguides (ZMWs) imaging surface via a biotin at the 3’end of the mRNAs (Fig. S1, A) (*1*). After washing away unbound components, 60S (Cy5-labeled), eIF5B (Cy5.5-labeled) and the first elongator Phe-tRNA^Phe^ (Cy3.5-labeled) in ternary complex with the elongation factor eEF1A and GTP (Phe-TC) were delivered along with required eIFs to start the reaction. By directly illuminating all the fluorescent dyes, we could monitor, in real time, the order of molecular events occurring during the late translation initiation stages and its transition to elongation (Fig. S1, B) (*1*). This allowed us to measure and determine the occupancy times of eIF5B on newly formed 80S complexes on the native reaction pathway. In particular, for the model mRNA-Kozak, the mean time of 60S joining to form an 80S was ~16 s with the eIF5B occupancy time on the 80S ~34 s at 20°C and 3 mM free Mg^2+^; whereas these values were ~3.6 s and 68.7 s, respectively, at 20°C and 10 mM free Mg^2+^ (*1*). Simulating the kinetic curves from these two reactions provided the information about the time-evolution of the populations of different complexes (Fig. S1, C) (*1*). To aid the reconstruction of a high resolution structure of the on-pathway eIF5B-bound 80S prior to its transition to elongation without the need of non-hydrolysable GTP analogs or mutants, we decided to prepare and freeze our sample at timepoint 40 s at 20°C and 10 mM free Mg^2+^, where we would expect the eIF5B-bound 80S population accounts ~60% of the total 80S particles (Fig. S1, C).

Samples were prepared in the buffer containing 30 mM HEPES-KOH pH 7.5, 100 mM KOAc, 10 mM Mg(OAc)_2_ and 1 mM GTP:Mg^2+^. First, a ternary complex mixture was prepared by pre-incubating 3.8 μM eIF2 at 30°C for 10 min, followed by another 5 min incubation at 30°C after addition of 2.8 μM Met-tRNA_i_. Next, this ternary complex mixture was diluted to one third of the concentration, and incubated at 30°C together with 1 μM eIF1, 1 μM eIF1A, 0.5 μM model mRNA and 0.3 μM 40S for 15 min, resulting a 48S PIC mixture. Separately, a 60S mixture was prepared by mixing 1 μM eIF5, 1μM eIF5B and 0.15 μM 60S. The 48S PIC and 60S mixtures were kept on ice before sample freezing. In parallel, the 200-mesh Quantifoil R2/1 grids (Electron Microscopy Sciences, Q250AR1) were glow-discharged for 25 s in a PELCO easiGlow glow discharger (Ted Pella, Inc.). After prewarming the samples to room temperature, 3.5 μL of the 48S PIC mixture was mixed with 3.5 μL of the 60S mixture. A 3 μL sample from the resulted mixture was applied to each grid at 21 °C and 95% humidity. The sample was vitrified by plunging into liquid ethane after 2.5 s blotting using a Leica EM GP (Leica Microsystems) plunger. Total time from combining the 48S PIC and 60S mixtures to grid freezing was ~40 s.

#### Single-molecule experiments comparing Met-tRNA_i_^Met^ and Gly-tRNA_i_^Met^

Native yeast tRNA_i_^Met^ was mis-acylated by the flexizyme dFx with Gly-DBE as described (*2, 3*). Real-time single-molecule experiments on the ZMW-based PacBio RSII instrumentation and data analyses were performed with exactly the same methodology as previously described (*1*). The mean times of 60S arrival to the immobilized 48S PICs (60S arrival time) and the subsequent transition to elongation (*Δ*t, Fig.S1, **B**) were determined in experiments performed with the model mRNA-Kozak and unlabeled eIF5B at 3 mM free Mg^2+^ and 20°C.

#### Generation of eIF5B mutant and growth experiments in yeast

The *Saccharomyces cerevisiae fun12Δ* strain J111 (*MATa ura3-52 leu2-3 leu2-112 fun12Δ*) (*2*) lacking eIF5B was transformed with an empty vector (eIF5BΔ) or plasmids expressing N-terminally deleted (lacking residues 28-396) form of wild-type (WT) eIF5B (*4*) or the eIF5B-R955A mutant, as indicated. Transformants were grown to saturation, and 5 µl of serial dilutions (of OD_600nm_ = 1.0, 0.1, 0.01, 0.001, and 0.0001) were spotted on minimal medium supplemented with essential nutrients (SD) or medium containing 10 mM 3-aminotriazole (3-AT) to cause histidine starvation. Plates were incubated 3 days at 30°C.

#### Image processing and structure determination

Contrast transfer function parameters were estimated using CTFIND4 (*5*) and particle picking was performed using Relion3.1(*6*) without the use of templates and with a diameter value of 260 Ångstrongs. All 2D and 3D classifications and refinements were performed using RELION (*6–8*). An initial 2D classification with a 4 times binned dataset identified all ribosome particles. A consensus reconstruction with all 80S particles was computed using the AutoRefine tool of RELION. Next, 3D classification without alignment and a mask including the inter-subunit space and the 40S head (four classes, T parameter 4) identified a class with unambiguous density for eIF5B and a tRNA in the P site (*7*). This class was independently processed with unbinned data, yielding high resolution maps with density features in agreement with the reported resolution. Local resolution was computed with RESMAP (*9*).

#### Model building and refinement

Models from yeast 40S, 60S (*10*), tRNA_i_ and eIF5B were docked into the maps using CHIMERA (*11*), and COOT (*12*) was used to manually adjust these initial models. An initial round of refinement was performed in Phenix using real-space refinement (*13*) with secondary structure restraints and a final step of reciprocal-space refinement with REFMAC (*14*). The fit of the model to the map over-fitting tests were performed following standard protocols in the field (*15*).

---

**Fig. S1.**
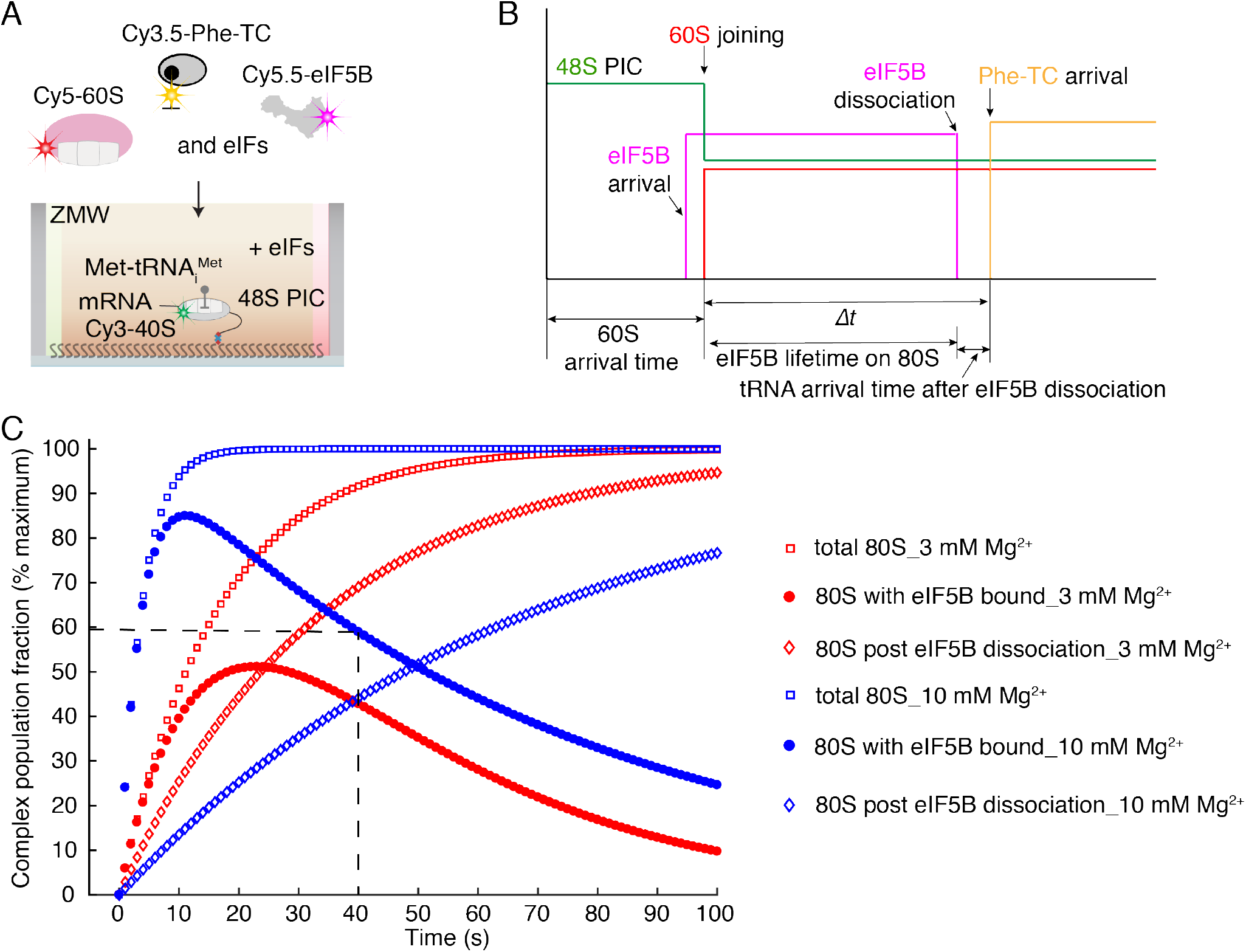
CryoEM sample preparation guided by single-molecule dynamics. (**A**) Single-molecule fluorescence microscopy experimental set up and (**B**) representative schematic experimental trace with the molecular events annotated (*1*). Mean times for subunit joining to form the 80S IC (60S arrival time) and subsequent transition into elongation (*Δ*t, signaled by the arrival of the first elongator Phe-TC) were estimated. *Δ*t is obtained by the addition of the eIF5B lifetime on 80S and the tRNA arrival time after eIF5B dissociation, with the latter being much smaller than the former. (**C**) Simulated time-evolution of 80S complex populations based on the previously determined kinetics in experiments performed with the model mRNA-Kozak at 20°C (*1*). Informed by our kinetics measurements, cryoEM samples corresponding to the timepoint 40 s after mixing 48S PICs, eIF5B:GTP and 60S in the presence of required eIFs at 10 mM free Mg^2+^ and 20°C were frozen. Dashed line indicates the estimated timepoint when the 80S population with eIF5B is ~60%.

**Fig. S2.**
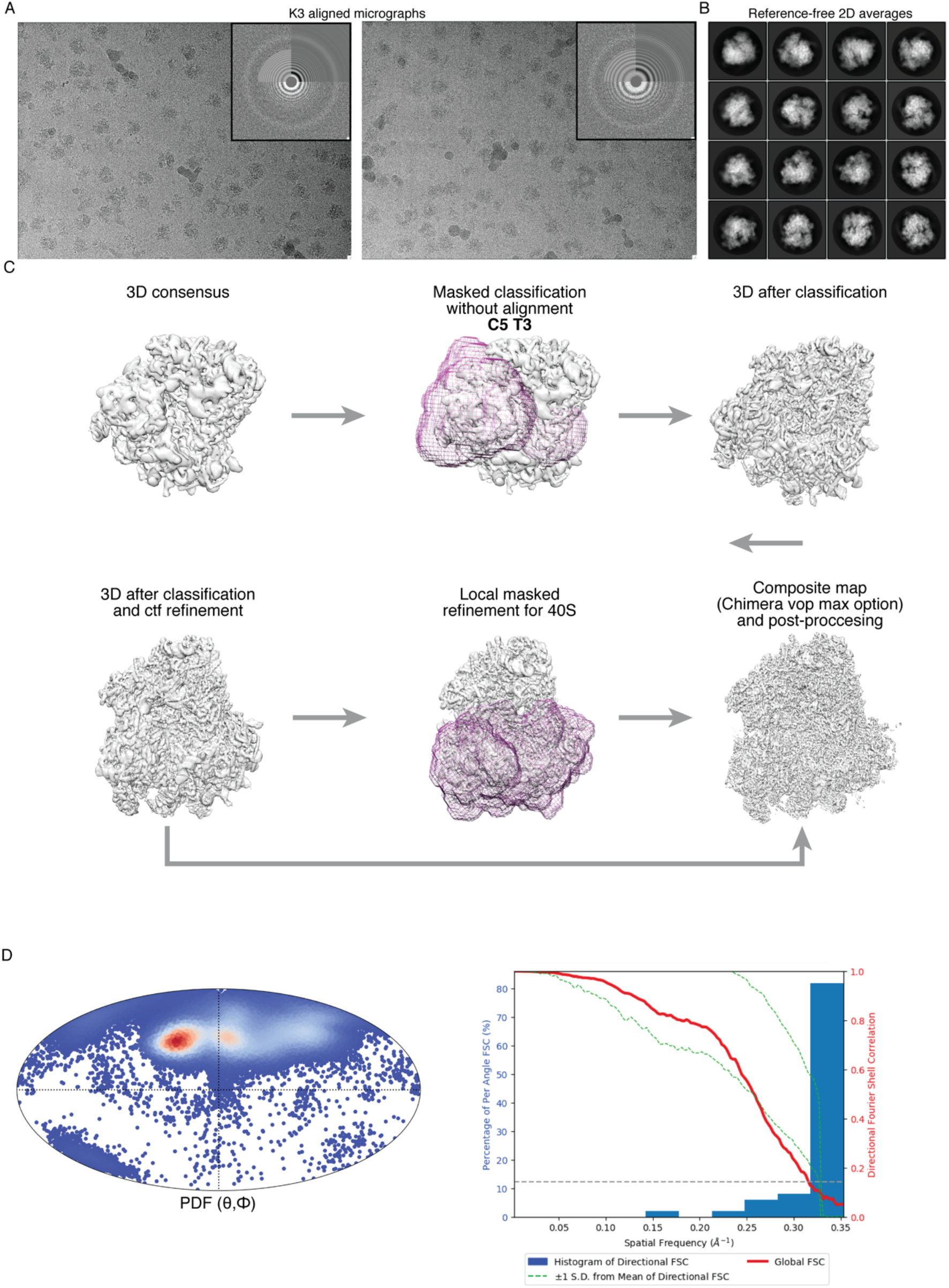
Representative cryoEM images, reference-free 2D averages and image processing workflow. **(A)** Motion corrected micrographs. (**B**) Selected 2D averages. (**C)** Classification workflow. (**D**) On the left, Eulerian angles distribution of the final class and on the right directional FSC (*16*).

**Fig. S3.**
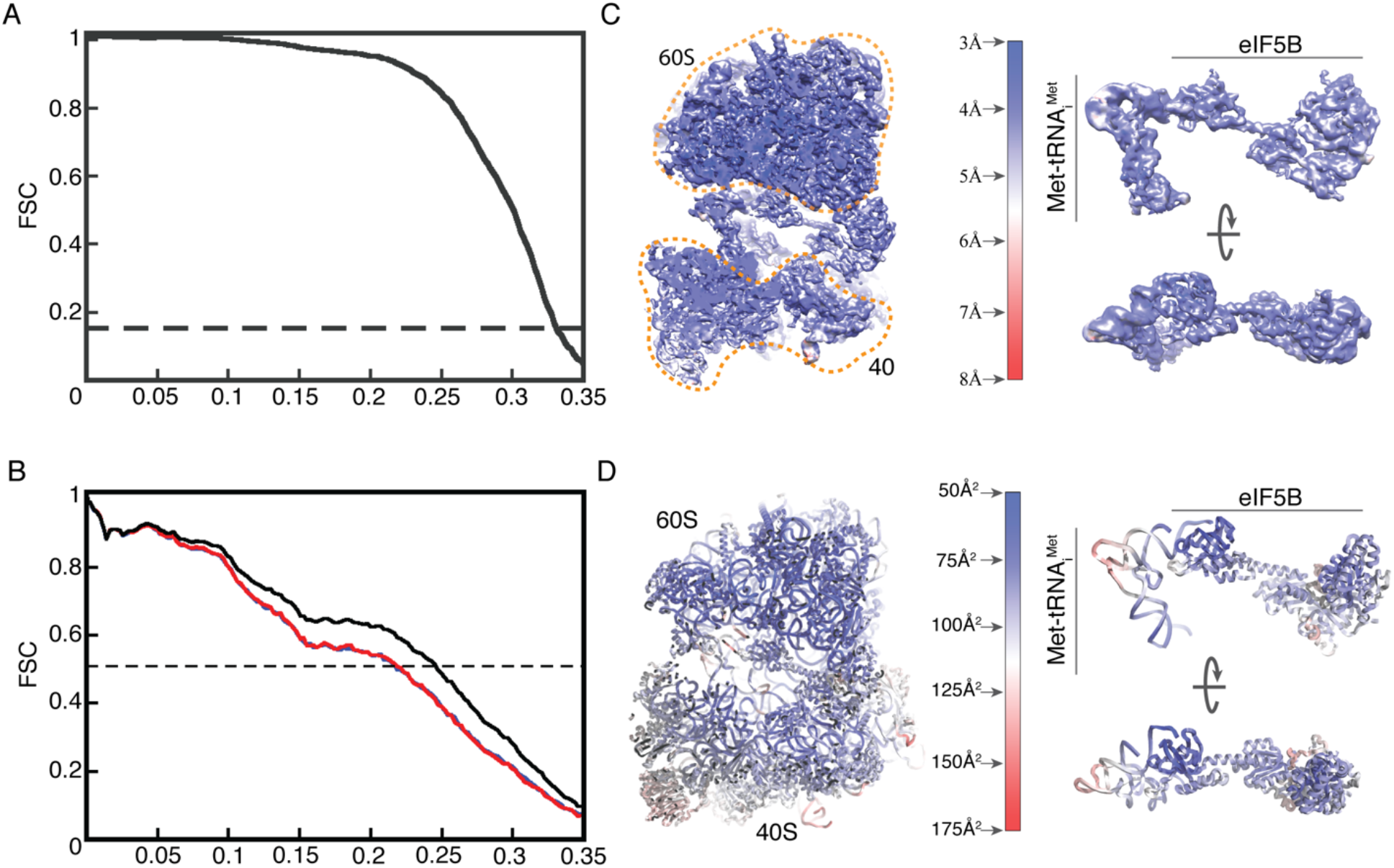
Fourier Shelf Correlation curves, model validation and local resolution. **(A)** Fourier Shell Correlation curve (FSC) for independently refined half-maps indicated a final resolution of 2.9Å. **(B)** Model validation FSC for the final “shacked” model and half-map 1 (blue line) and half-map 2 (red curve) not included in the refinement. Black line represents the FSC for the final model against the final, post-processed map used in model building and model refinement. The overlapping of the blue and red curve guarantees absence of overfitting in our model (*15*). **(C)** Unsharpened map colored according to local resolution calculations. On the right, a detailed view for the region of the map containing density for eIF5B and Met-tRNA_i_^Met^. **(D)** Final stereochemically refined model colored according to temperature (B) factors estimated by REFMAC (*14*). Regions of the model with higher B factors correlate with flexible areas of the map.

**Fig. S4.**
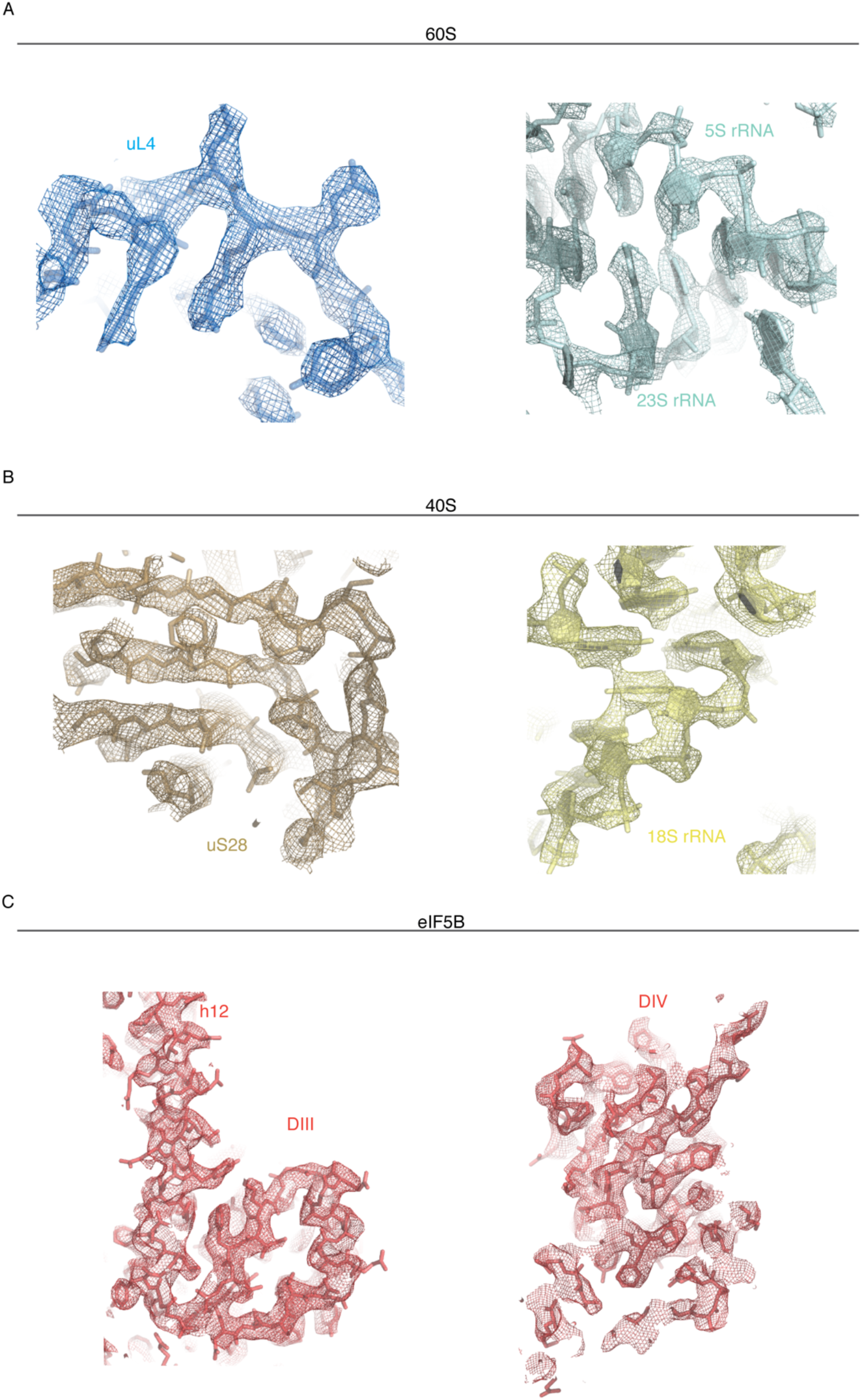
CryoEM representative densities. Final, post-processed cryoEM densities for: **(A)** 60S components, **(B)** 40S components and **(C)** eIF5B.

**Fig. S5.**
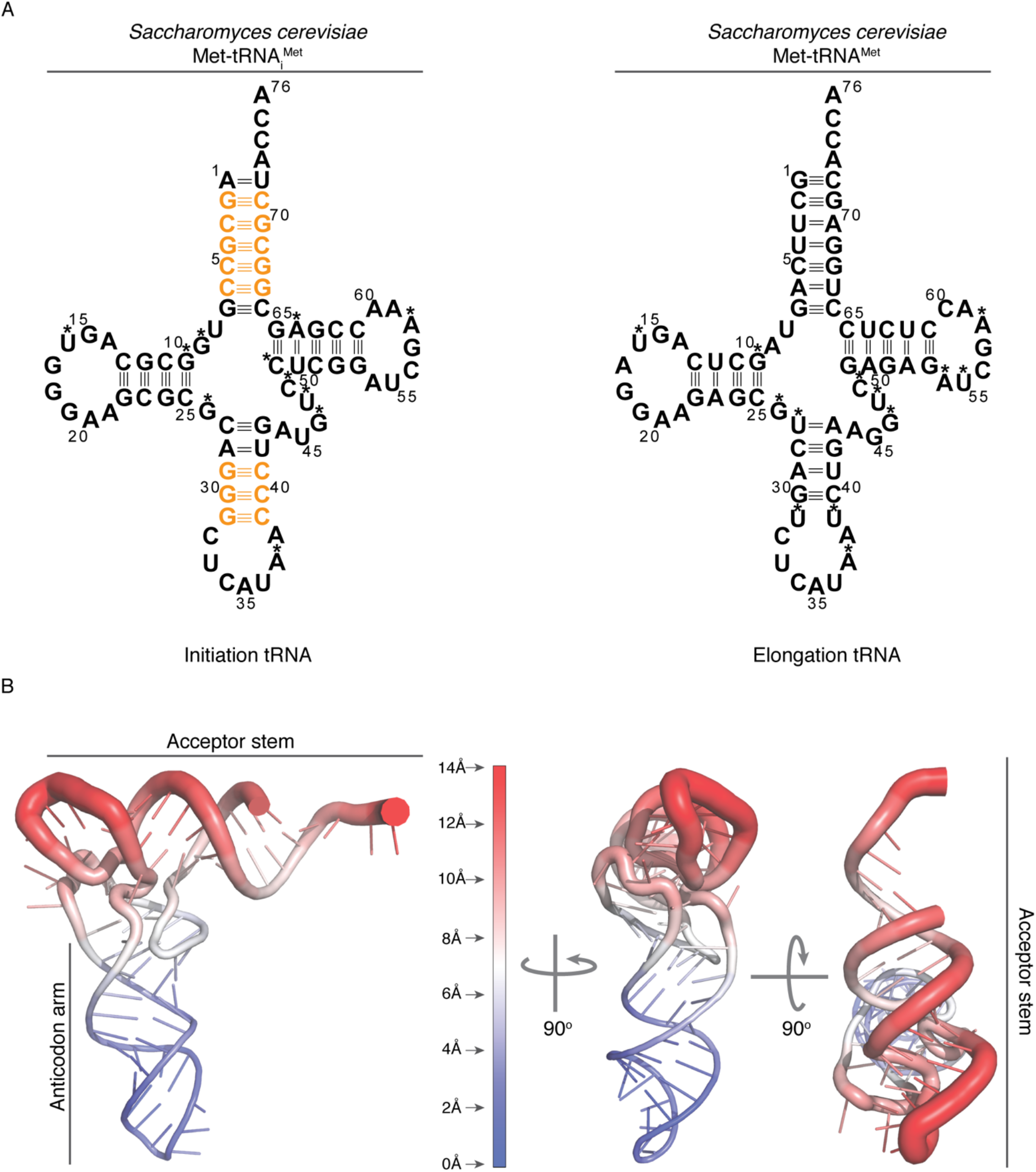
*Saccharomyces cerevisiae* tRNA^Met^ sequences. **(A)** Left, *S.cerevisiae* sequence for tRNA_i_ with the G-C base pair clusters unique to initiator tRNA in orange. Right, elongation tRNA^Met^ sequence in *S.cerevisiae.* Asterisks denote modified nucleotides. **(B)** Root mean square deviations (r.m.s.d) between atoms computed for tRNAi in an elongation configuration versus the configuration described in this work. Mayor differences are found within the acceptor stem.

**Fig. S6.**
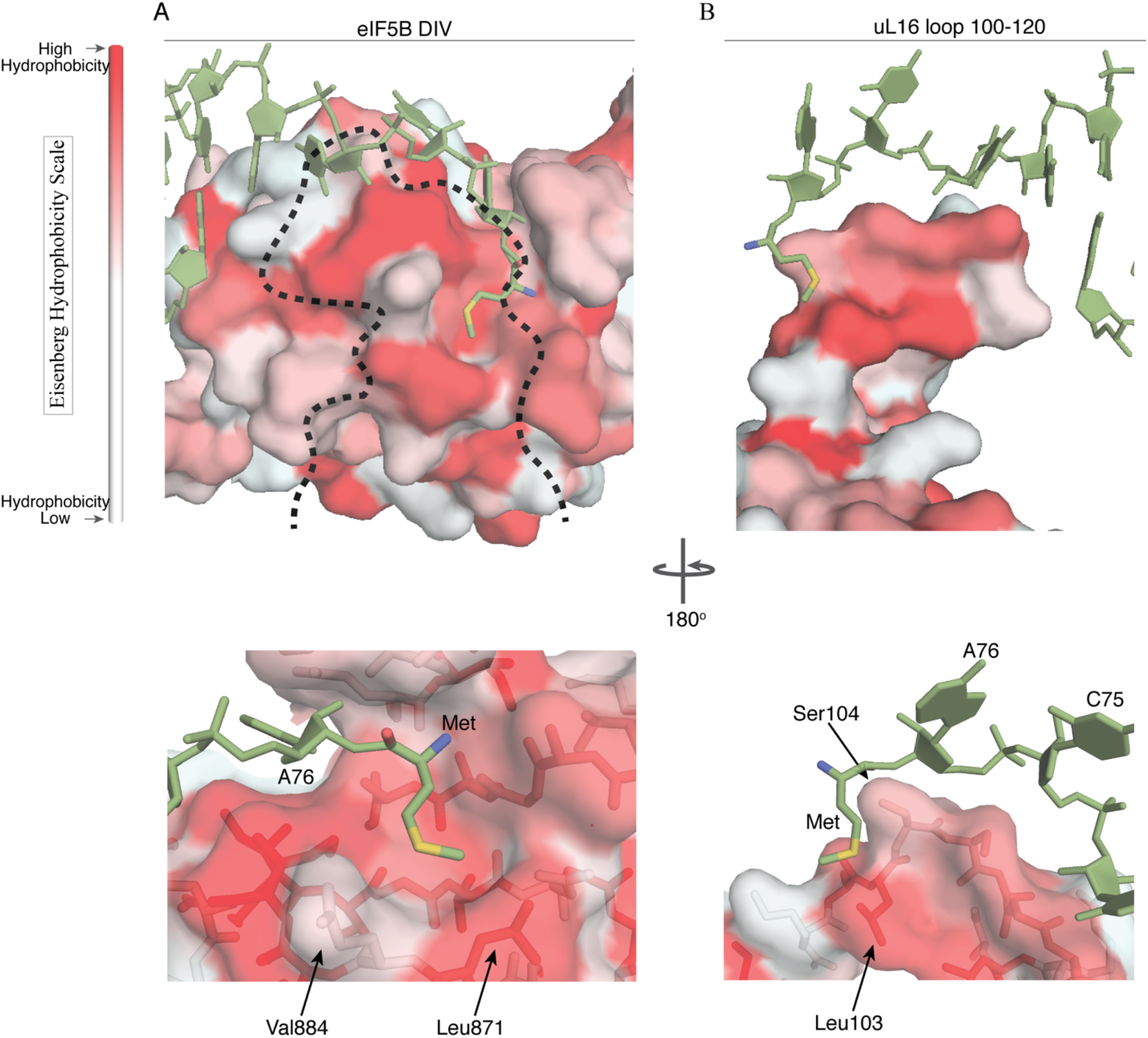
Surface hydrophobicity calculations for eIF5B and uL16. **(A)** eIF5B domain IV represented as Van der Waals surface colored according to the Eisenberg hydrophobicity scale. In sticks is shown Met-tRNA_i_^Met^ (*17*). Indicated with a dash line is the position occupied by uL16 loop 100-120. **(B)** Same representation as in **(A)** from the uL16 view. Methionine residue esterified to the 3’OH of Met-tRNA_i_^Met^ A76 is hosted in a highly hydrophobic cavity formed by eIF5B DIV and uL16 loop 100-120.

**Fig. S7.**
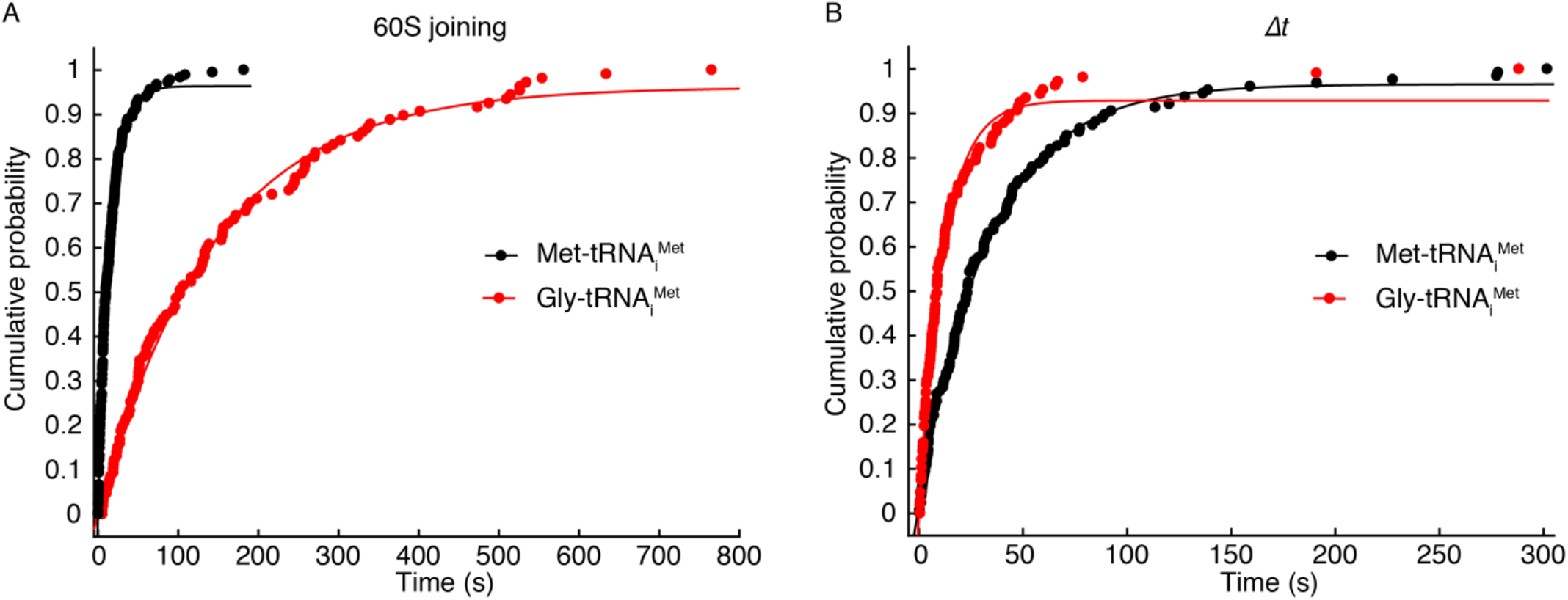
Kinetics of subunit joining and subsequent transition into elongation in experiments performed with Met-tRNA_i_^Met^ and Gly-tRNA_i_^Met^. The cumulative probability distributions of the dwell times for 60S joining **(A)** and the transition to elongation (*Δt*, **B**) from experiments performed at 3 mM free Mg^2+^ and 20 °C with the model mRNA-Kozak and dark eIF5B. Data points were fitted to a single-exponential equation. When Met-tRNA_i_^Met^ was used, the mean 60S arrival time was 14.8 s (± 0.4 s, 95% confidence interval), with *Δt* of 33.7 s (± 0.9 s, 95% confidence interval) (number of molecules analyzed = 127), similar to those values determined previously under the same experimental conditions (*1*). For Gly-tRNA_i_^Met^ (red) the mean 60S arrival time increased to 142.9 s (± 6.2 s, 95% confidence interval), and *Δt* decreased to 11.7 s (± 0.5 s, 95% confidence interval) (number of molecules analyzed = 107).

**Fig. S8.**
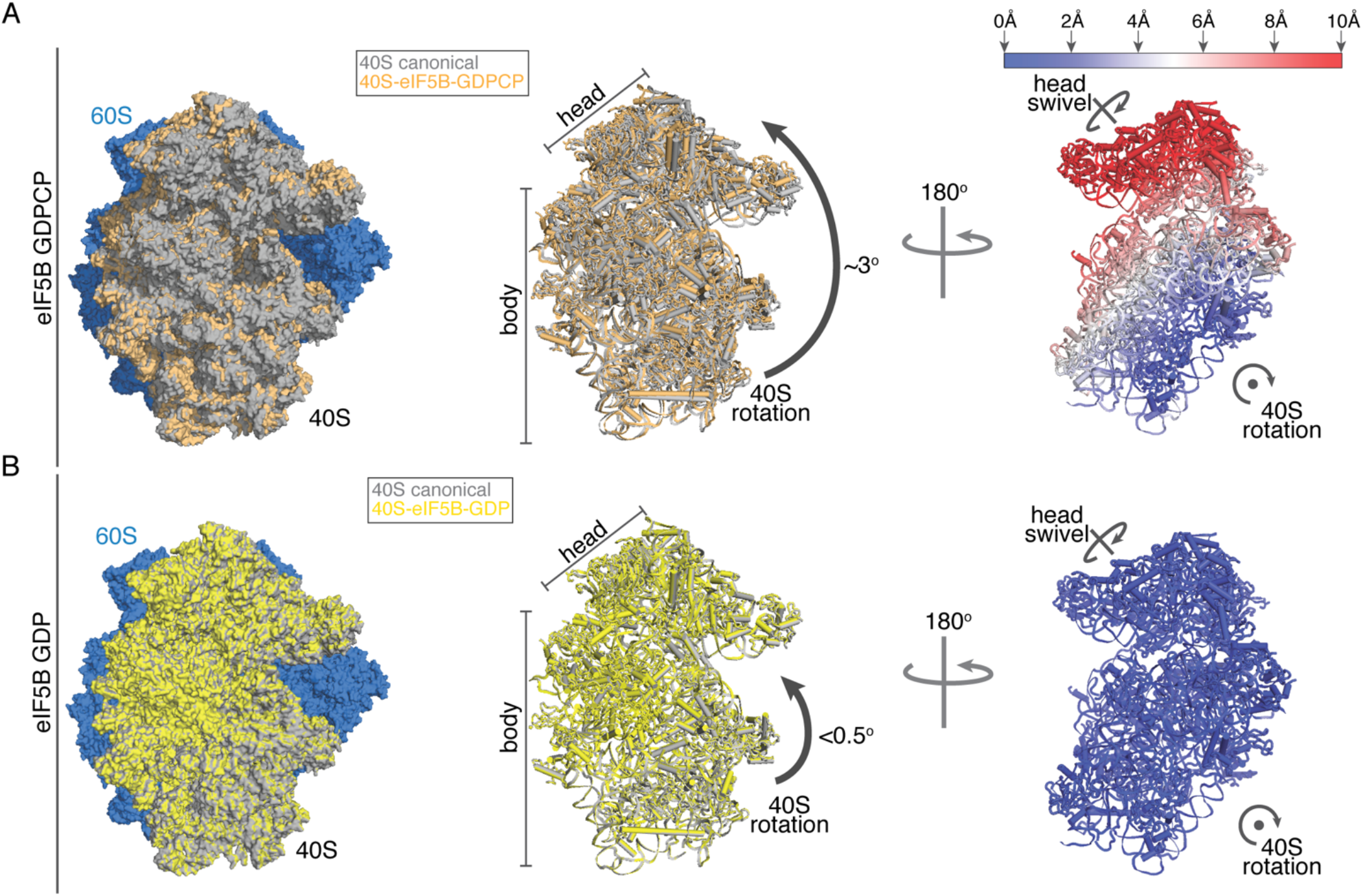
40S subunit rotation state in eIF5B containing complexes. **(A)** In the presence of a non-hydrolysable GTP analog, binding of eIF5B to the 80S induces a moderate degree of 40S rotation compared with a canonical state (*10, 18*). **(B)** In the present reconstruction, with eIF5B in the 80S complex and hydrolyzed GTP, the 40S subunit features an almost canonical configuration with very little rotation and/or head swivel.

**Movie S1. Met-tRNA_i_^Met^ transition from late initiation to elongation.**

The interaction DIV of eIF5B establishes with the acceptor stem of Met-tRNA_i_^Met^ keep the _73_ACCA_76_-Met of the tRNA away from the PTC (blue). Upon eIF5B departure, the _73_ACCA_76_-Met is free to accommodate in the PTC so the Met-tRNA_i_^Met^ adopts a full elongation-competent conformation.

**Table S1.**
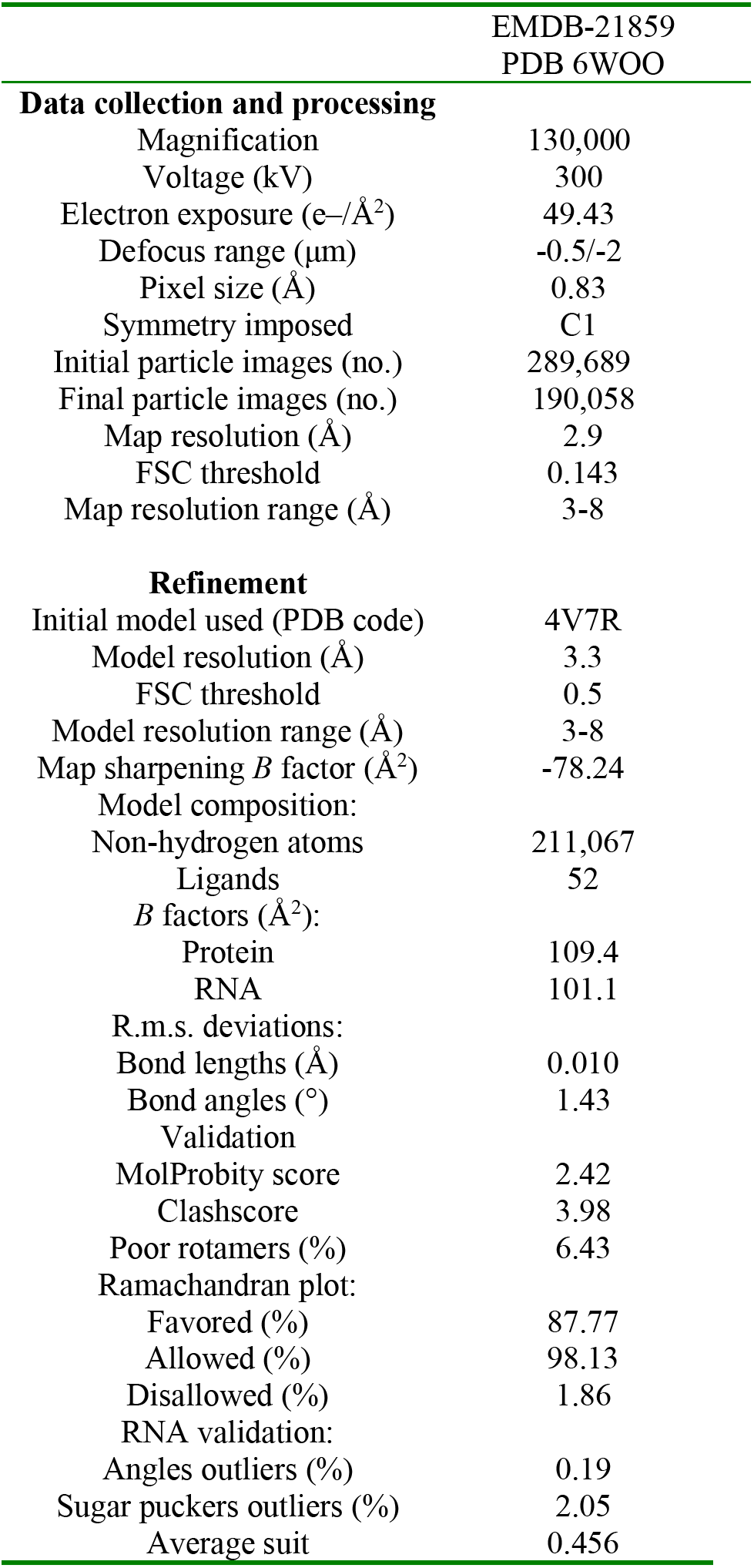
Cryo-EM data collection, refinement and validation statistics.

